# Mitochondrial dysfunction underlies impaired neurovascular coupling following traumatic brain injury

**DOI:** 10.1101/2023.07.20.549872

**Authors:** Gerben van Hameren, Jamil Muradov, Anna Minarik, Refat Aboghazleh, Sophie Orr, Shayna Cort, Keiran Andrews, Caitlin McKenna, Nga Thy Pham, Mark A. MacLean, Alon Friedman

## Abstract

Traumatic brain injury (TBI) involves an acute injury (primary damage), which may evolve in the hours to days after impact (secondary damage). Seizures and cortical spreading depolarization (CSD) are metabolically demanding processes that may worsen secondary brain injury. Metabolic stress has been associated with mitochondrial dysfunction, including impaired calcium homeostasis, reduced ATP production, and elevated ROS production. However, the association between mitochondrial impairment and vascular function after TBI is poorly understood. Here, we explored this association using a rodent closed head injury model. CSD resulted in neurobehavioral decline after TBI. Craniotomy was performed to elicit CSD via electrical stimulation or to induce seizures via 4-aminopyridine application. We measured vascular dysfunction following CSDs and seizures in TBI animals using laser doppler flowmetry. We observed a more profound reduction in local cortical blood flow in TBI animals compared to healthy controls. Following TBI, CSD resulted in mitochondrial dysfunction and pathological signs of increased oxidative stress adjacent to the vasculature. We explored these findings further using electron microscopy and found that TBI and CSDs resulted in vascular morphological changes and mitochondrial cristae damage in astrocytes, pericytes and endothelial cells. Overall, we provide evidence that CSDs induce mitochondrial dysfunction, impaired cortical blood flow, and neurobehavioral deficits in the setting of TBI.

**Highlights:**

Cortical spreading depolarization after TBI causes behavioral decline in rats.
Vasoconstriction and oligemia after cortical spreading depolarization is worse in TBI brains.
Spreading depolarization causes impaired mitochondrial function.
TBI and spreading depolarization result in constricted vessels and increased pericyte size.
TBI and spreading depolarization result in mitochondrial damage in vascular cells.

**Graphical abstract:** 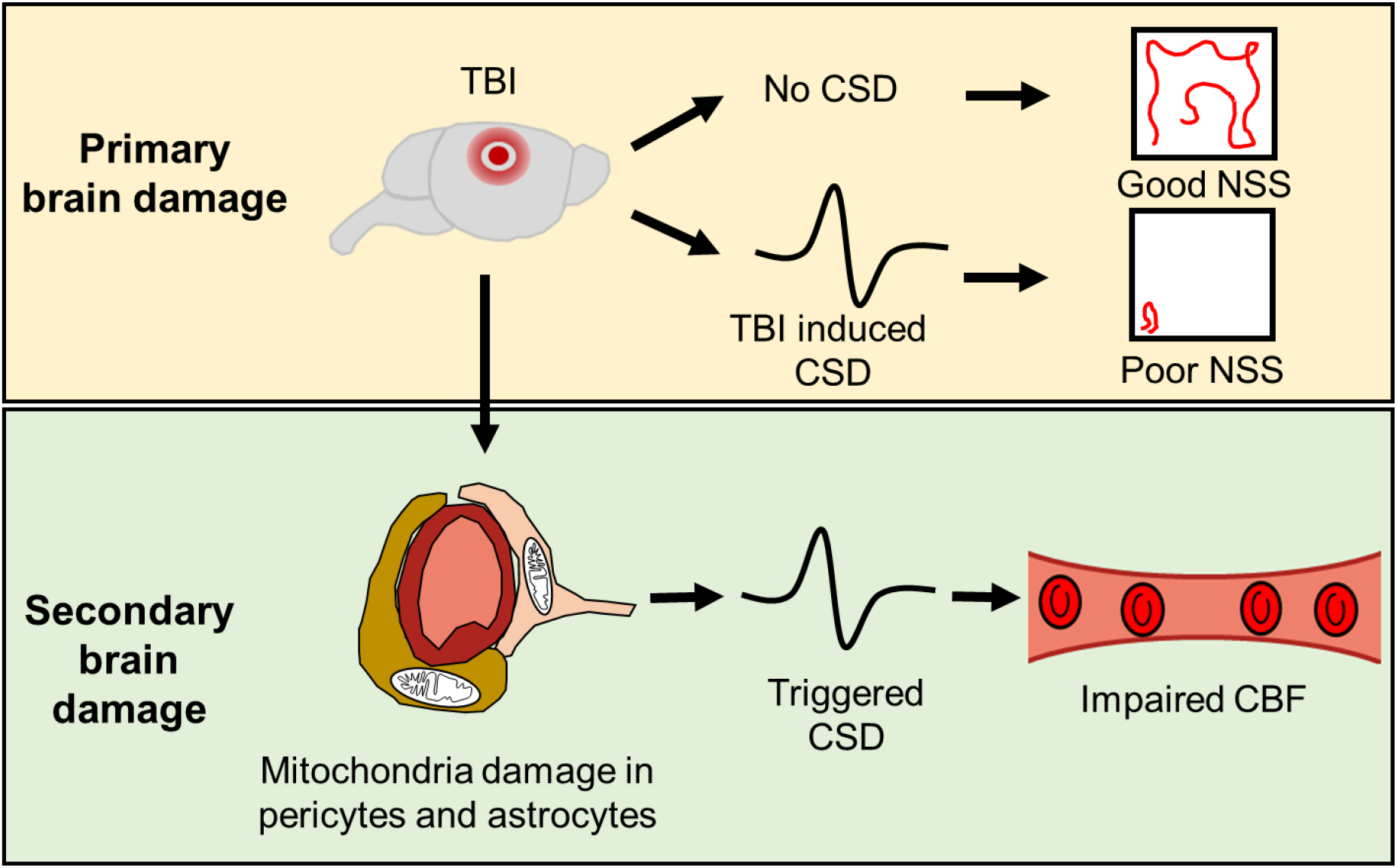

## 1. Introduction

Traumatic brain injury (TBI) is a major global source of morbidity and mortality, affecting 40 million people annually worldwide (Haarbauer-Krupa et al., 2021). TBI-related injury is classified as primary or secondary (Shaikh and Waseem, 2022). Primary injury occurs instantaneously and may involve cerebral contusions, intracranial hematomas, white matter tract sheering, and disruption of the brain cellular integrity. In contrast, secondary injury develops in the minutes, hours, or days after impact. Mechanisms underlying secondary injury are complex and not well understood.

The electrophysiological changes that occur within minutes after head impact may mechanistically trigger secondary brain damage processes (Hartings et al., 2020). Following head impact, neurons are depolarized to generate cortical spreading depolarization (CSD) or, rarely seizures (Aboghazleh et al., 2021). CSD is a propagating wave of synchronous depolarization of a large set of neuronal populations (Dreier, 2011; Dreier et al., 2018, 2012; Hartings et al., 2011; von Baumgarten et al., 2008). CSD has been described as the most common (50-80%) pathological electrophysiological process after TBI in both animal studies and human recordings (Aboghazleh et al., 2021; Fabricius et al., 2006; Parker et al., 2022; Strong et al., 2002). The net influx of cations during CSD causes a near-complete sustained neuronal depolarization, which blocks further initiation of action potentials (spreading depression of neuronal activity) (Dreier, 2011; Dreier et al., 2019, 2018). Seizures are similarly characterized by depolarization of larger neuronal population, but with excessive firing of action potentials (March, 1998).

Restoration of ion gradients following CSD or seizures is essential for repolarization and regaining of physiological neuronal activity. The need of excess adenosine triphosphate (ATP) for sodium/potassium ATPase functioning therefore requires adaptation of mitochondrial function (Toglia and Ullah, 2019) . In addition to ATP production via oxidative phosphorylation, mitochondria produce reactive oxygen species (ROS) and buffer intracellular calcium (Brookes et al., 2004). The uptake of Ca^2+^ ions during CSD is associated with depolarization of the inner mitochondrial matrix (Bahar et al., 2000; Zhou et al., 2010) and influx of water, leading to mitochondrial swelling and cristae damage (Kirov et al., 2020). However, the exact role of mitochondria following TBI and CSD remains unknown.

In addition to neurons, the function of astrocytes, pericytes, and endothelial cells within the neurovascular unit (NVU) can become impaired by CSD (Khennouf et al., 2018; Risher et al., 2012; Sadeghian et al., 2018; Sword et al., 2013). Pericytes are contractile cells that regulate cerebral blood flow (CBF) in capillaries in response to neurotransmitters, systemic oxygen levels, ATP availability, nitric oxide, and intracellular calcium levels (Attwell et al., 2016; Burdyga and Borysova, 2014; Hall et al., 2014). Reduced mitochondrial respiration and ATP production has also been associated with reduced contractile capability of pericytes (Liu et al., 2021). Astrocytes regulate CBF via interactions between their endfeet and endothelial cells. Reduced CBF triggers calcium influx into astrocytes (Marina et al., 2020), which is involved in regulation of both vasoconstriction and vasodilation states (Gordon et al., 2008; Haidey and Gordon, 2021; Menyhárt et al., 2018). Therefore, TBI- and CSD induced mitochondrial dysfunction may be linked with vascular morphological and CBF changes, but this remains to be determined

We hypothesized that mitochondrial damage may underlie impaired vascular function and contribute to secondary injury following TBI, particularly in the context of metabolically demanding electrophysiological events such as CSD and seizures (Hiebert et al., 2015). As such, our overarching study objective was to test the association between mitochondrial damage and vascular dysfunction under metabolically challenging conditions (i.e. CSD and seizures) using a rodent model of TBI.

## 2. Methods

### 2.1 Animals

Experiments were performed following institutional approval (protocols: 20-014 and 17-105 and 21-103) by the Dalhousie University Committee on Laboratory Animals. Experiments were conducted in accordance with the Canadian Council on Animal Care.

### 2.2 Traumatic Brain Injury

Single moderate TBI was induced using a modified weight-drop model (Fig. 1A, Marmarou et al., 1994; Mychasiuk et al., 2014; Parker et al., 2022). Adult, 9-12 week old, male Sprague Dawley rats (n=79, Table 1) were sedated using an induction chamber (3% isoflurane, 2 L/min O_2_), until the toe-pinch reflex was absent. Rats were then placed in the prone position on a sheet of aluminum foil taped to the top of a plastic box (30x30x20 cm in depth). A metal bolt (1 cm diameter x 10 cm length) was placed on the rat’s head anterior to the lambda suture line (but posterior to bregma) in the midline via alignment with the animal’s ears as an anatomical reference. Moderate TBI was induced using a weight (450g) travelling vertically for 1.02 m along a metal guide rail, impacting the metal bolt, transmitting the energy onto the rat’s head. Following the impact, animals fell through the foil onto a foam pad placed at the bottom of the box, causing a rotation of the head and neck. Immediately upon impact, a peripheral oxygen saturation (spO_2_) sensor (ML325, ADInstruments) was placed on the right hind paw, and recorded using and a PowerLab data acquisition device (PL3508, ADInstruments) and LabChart 8 software (AD Instruments). Control animals were anesthetized but did not receive TBI. Two days after TBI or sham anesthesia, animals were used in follow-up experiments (see below) or sacrificed, and skull and brains were photographed for anatomical examination.

**Table 1.** Number of animals per group and respective measurements.

|  | Total | spO<sub>2</sub> | NSS | anatomy | ATPase | ECoG recording | brightfield imaging | LDF | cortical O<sub>2</sub> | mitoSOX | Rhod2-AM | electron microscopy |
| --- | --- | --- | --- | --- | --- | --- | --- | --- | --- | --- | --- | --- |
| **<u>Isoflurane control</u>** | **<u>103</u>** |  |  |  |  |  |  |  |  |  |  |  |
| No surgical intervention | 11 | 7 |  | 4 | 4 |  |  |  |  |  |  | 4 |
| **<u>1.5x5 mm craniotomy</u>** | **<u>78</u>** |  |  |  |  |  |  |  |  |  |  |  |
| CSD induced | 64 |  |  |  |  | 25 | 16 | 17 | 11 | 13 | 10 | 5 |
| seizures induced | 10 |  |  |  |  | 7 |  | 7 | 2 | 3 |  |  |
| sham surgery | 4 |  |  |  |  | 10 |  |  |  |  |  | 4 |
| **<u>1x1 mm craniotomy</u>** | **<u>14</u>** |  |  |  |  |  |  |  |  |  |  |  |
| CSD induced | 6 |  | 6 |  |  | 6 |  |  |  |  |  |  |
| sham surgery | 8 |  | 8 |  |  | 8 |  |  |  |  |  |  |
| **<u>TBI without ECoG implant</u>** | **<u>79</u>** |  |  |  |  |  |  |  |  |  |  |  |
| No surgical intervention | 30 | 9 | 4 | 21 | 4 |  |  |  |  |  |  | 4 |
| **<u>1.5x5 mm craniotomy</u>** | **<u>49</u>** |  |  |  |  |  |  |  |  |  |  |  |
| CSD induced | 42 |  | 26 |  |  | 20 | 13 | 17 | 9 | 13 | 8 | 4 |
| seizures induced | 7 |  |  |  |  | 7 |  | 7 |  | 7 |  |  |
| **<u>TBI with ECoG implant</u>** | **<u>71</u>** | 3 | 18 |  |  | 71 |  |  |  |  |  |  |

### 2.3 Neurological severity scoring (NSS)

NSS testing was performed, as described previously (Parker et al., 2022), via a composite of open field, beamwalk and inverted mesh test scores (four points each).

### 2.4 Na^+^/K^+^ ATPase assay

Two days after TBI, animals were sacrificed and a subset of brains (4 TBI brains, 4 controls) were snap frozen in liquid nitrogen. Obtained brains were then dissected, placed on ice and the cortex was isolated and homogenized in 10 ml homogenization solution (0.25 M sucrose, 1.25 mM EGTA, and 10 mM Tris buffer) per 1 g wet weight. Na^+^/K^+^ ATPase activity from brain homogenates was tested using a spectrophotometric enzyme kinetic assay (Reiffurth et al., 2020). In this assay, the rate of NADH oxidation is measured, which is dependent on the amount of ATPase in the dissected tissue. 10 µM of ouabain, a Na^+^/K^+^ ATPase inhibitor was added, to test the α2/3 Na^+^/K^+^ ATPase isoforms selectively. Each sample was tested in triplicate.

### 2.5 Electrocorticography (ECoG) electrode implants

A separate cohort of animals (n=71) was implanted with ECoG electrodes 24 to 72 hours prior to TBI under deep isoflurane anesthesia (3% for induction and 1.5% for maintenance) as described earlier (Aboghazleh et al., 2021). SpO_2_ was monitored continuously (ML325, ADInstruments). A midsagittal incision (2.5 cm) was made to expose the skull and two holes (2x2 mm in diameter) were drilled for screw placement (stainless steel bone screws, Fine Science Tools, 0.86 mm x 4 mm) in both parietal bones (2 mm posterior to bregma, 2 mm anterior to lambda, and 3 mm lateral to sagittal suture). A ground electrode was inserted into neck subcutaneous tissue. Epidural ECoG electrodes were constructed from Teflon insulated silver wire (280 µm diameter, A-M Systems, Inc.) and miniature connectors (Ginder Scientific, ABS plug, GS09PLG-220). The silver wires of the electrodes were wrapped around the screws and were fixed to the skull using dental cement. A cylindrical platform (1 cm in diameter and 1.5 cm in height) was formed above the frontal or parietal bones using dental cement, via which the dropped weight was used to transmit an impact to the brain. Animals were tethered to an Octal bio amplifier (ML138, ADInstruments, Sydney, Australia) for recording differential epidural ECoG signals. Near-direct current (DC) recordings were acquired (sampling rate of 1 kHz) with a high-pass filter of 0.02 Hz, a low-pass filter of 100 Hz, and a 60 Hz notch filter. Alternative current (AC) recordings were obtained by applying a 0.5-45 Hz bandpass to the near-DC-ECoG recording. A one-hour baseline recording was acquired in a recording box (60x30x30 cm). Immediately following TBI, animals were reconnected to the recording system within < 30 seconds. Electrical recordings were continued for 2 hours following TBI. ECoG data were analyzed using LabChart software (version 8) and MATLAB.

### 2.6 Craniotomy, electrocorticography and CBF measurement in vivo

Rats (n=127) underwent craniotomy under anesthesia (1.5-3% isoflurane) and cortical vascular imaging was performed via an open window technique (Aboghazleh et al., 2022; Schoknecht et al., 2021). A cohort of animals (n=49) received TBI two days prior to craniotomy. In short, 10 µL of bupivacaine (Sterimax, ref. 02443694) and Vetergesic Multidose (0.1 mg/kg, Ceva Animal Health, ref. 02342510) were administered subcutaneously. A parietal cranial window (1.5 by 5 mm) was made, the dura was removed and artificial cerebrospinal fluid (aCSF) (129 mM NaCl, 21 mM NaHCO_3_, 1.25 mM NaH_2_PO_4_, 1.8 mM MgSO_4_, 1.6 mM CaCl_2_, 3mM KCl and 10 mM of glucose, pH 7.4) was applied to the exposed brain. A separate right frontal burr hole (1x1 mm) was also made. An Octal bio amplifier (ML138, ADInstruments, Sydney, Australia) was used for recording differential epidural ECoG signals via silver electrodes (ref. 786500, A-M Systems) placed on the cortex. A reference electrode was placed at an incision in the rat’s neck. Near-DC and AC recordings were obtained as described above. A Laser Doppler Flowmeter (AD Instruments) was immersed in aCSF over the parietal window to measure CBF velocity. For measurements of oxygen partial pressure, a probe (Unisense, OX-10) was inserted 1 mm into the cortex. ECoG recordings, local CBF and oxygen pressure were recorded in real time using LabChart 8 software (AD Instruments) or UniAmp software (Unisense, Sensortrace), respectively.

### 2.7 Induction of spreading depolarizations or seizures

Two stainless steel stimulation electrodes (0.5 mm in diameter and 2.5 cm in length) were placed at the right frontal bone window for epidural stimulation. CSDs were induced using electrical stimulation with 20 volt and 20 hertz for 2 seconds with a pulse duration of 5-20 ms (n=106 rats) (Bogdanov et al., 2016; Lückl et al., 2018). Repetitive CSDs were induced following a 15-20 minute interval. Seizures were induced by topical application of 100 µM 4-aminopyridine in aCSF (n=17 rats) applied directly to the cortical window.

### 2.8 NSS assessment after electrically triggered CSD

In a cohort of rats (n=14), a similar cranial window surgery was performed, but the size of the craniotomy was smaller (1x1 mm) without durotomy or placement of recording electrodes. CSD was electrically triggered via a second frontal burrhole, as described above. Occurrence of CSD was confirmed via changes in intrinsic optical signal through the parietal window. Five minutes after stimulation, the wound was sutured, after which the rat was moved to a recovery cage. Recovery from anesthesia was monitored and NSS was tested at twenty-five minutes following stimulation, using open field test, beamwalk test and inverted mesh test.

### 2.9 Assessment of mitochondrial function in vivo

Following craniotomy and removal of the dura (Section 2.6), MitoSOX Red (5 µM; ThermoFisher, ref. M36008) or Rhod2-AM (2 µM; Invitrogen, Ref. R1245MP) in aCSF was applied topically for 30 minutes with no ambient light for mitochondrial function assessment. The exposed parietal cortex was imaged (Axio Zoom V16, Zeiss GmbH and CMOS camera, PCO Edge5.5 model, PCO-Tech Canada) and dye fluorescence was visualized using 568/645 nm ex/em.

### 2.10 Electron microscopy

A subset of rats (n=21) was anaesthetized with sodium pentobarbital (100 mg/kg i.p.) at two days after TBI or within 1 hour following the last CSD and perfused transcardially with physiological saline and 2.5% glutaraldehyde solution. Brain samples were collected from six distinct cortical regions of both hemispheres. Tissue was rinsed with 0.1M sodium cacodylate buffer and post-fixed with 1.0% osmium tetroxide. Tissues were placed in 0.25% uranyl acetate at 4 °C overnight and then dehydrated in increasing concentrations of acetone (50, 70, 95, 100%). Tissues were then infiltrated with epon araldite resin. Finally, the tissue was embedded in pure epon araldite resin and heated at 60 °C in an oven for 48 hours. Thin sections (∼ 80-100nm thick) from cortical layers 1 and 2 were cut with a Diatome diamond knife using a Reichert-Jung Ultracut E Ultramicrotome, placed on 300 mesh copper grinds, and stained with uranyl acetate (2.0%) and lead citrate. Thin sections were analyzed for ultrastructural damage on a JEOL JEM 1230 Transmission Electron Microscope at 80kV. Images were captured using a Hamamatsu ORCA-HR digital camera.

### 2.11 Analysis and statistics

Quantitative analysis of electron micrographs was done using ImageJ (National Institutes of Health). Vessel circularity was measured by setting a region of interest around the lumen and calculated via the *Shape Descriptors* function. The average tight junction diameter was calculated by dividing total area of the tight junction by its length. Mitochondrial size was measured by a region of interest around the outer mitochondrial membrane. For cristae damage assessment, a threshold is created to distinguish between inner mitochondrial matrix and cristae, and threshold fraction is measured. Cell type identification was performed using morphological features, including electron density (Nahirney and Tremblay, 2021). Vascular diameter was measured using an in-house script in MATLAB as described previously (Levi et al., 2012). For *in vivo* fluorescent intensity measurements, multiple regions of interest were obtained in ImageJ and mean intensity of each region was calculated over the course of the video. Obtained data were corrected for background fluorescence and photobleaching. Statistical analysis was done using Graphpad Prism 9 (GraphPad Software, San Diego, California USA). Anderson-Darling tests were used for assessment of normal distribution. Significance was tested using Student t-tests, Mann-Whitney tests, Wilcoxon matched-pairs signed rank tests, one-way ANOVA with Kruskal-Wallis post-hoc test or two-way ANOVA with Sidak’s multiple comparison test. Statistical tests and p-values are indicated in the figure captions.

## 3. Results

### 3.1 Traumatic brain injury results in neurobehavioral decline and structural and functional changes

We first assessed functional, anatomical, and behavioral outcomes following a weight drop moderate TBI model (Fig. 1A). We measured a drop in spO_2_ acutely upon impact (Fig. 1B). The peak minimum saturation was observed 30 to 60 seconds post impact. Within 2 minutes, spO_2_ returned to levels indistinguishable to sham controls. Behavioral analysis showed reduced NSS scores (Fig. 1C) at 20 minutes after head impacts (Fig. 1C). At 48 hours following impact, the distribution of behavioral scores was bimodal (Fig. 1C). Post-mortem observation revealed bruises on the rat’s skull (Fig. 1D) and epidural hematomas with no evidence for subdural or cortical intracerebral bleeding (16 out of 21 animals, Fig. 1D). Sub-arachnoid haemorrhage was observed, most often adjacent the skull base region and the brainstem (15 out of 21, Fig. 1D). In addition, total ATPase activity in homogenates of cortical brain tissue was not significantly different between TBI animals and controls (Fig. 1E). Activity of the neuroglial α2/3 isoform ATPase activity was higher in TBI animals compared with controls (see methods) (Fig. 1E). The change in α2/3 isoform ATPase activity was measured 2 days after head impact, suggesting functional changes indicative of delayed, secondary brain injury.

**Figure 1.**
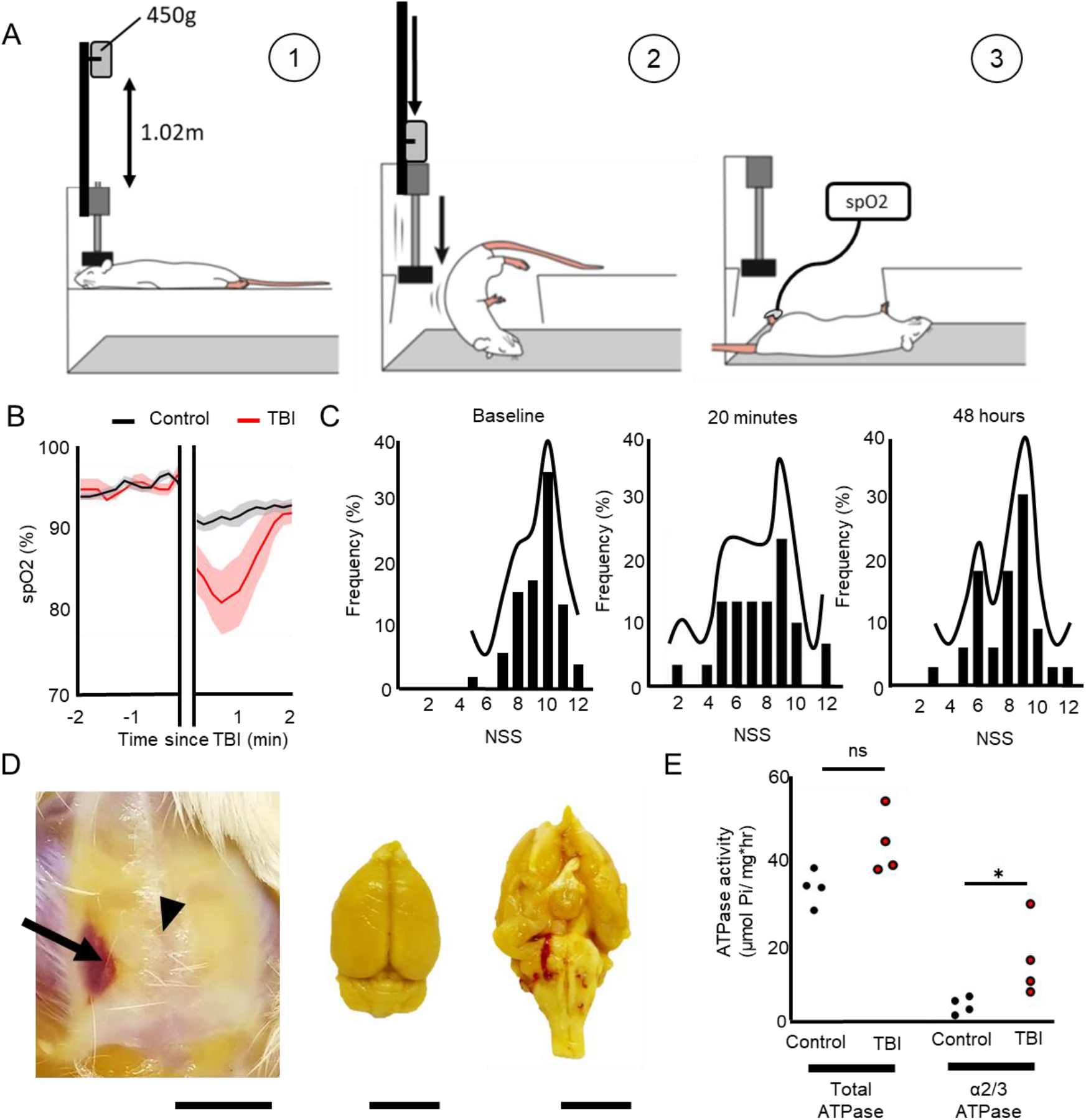
Single moderate TBI results in structural, functional and behavioral changes. **A)** An overview of the moderate TBI model. A weight of 450g is dropped from 1.02m, relaying a force on the head of an anesthetized rat that is lying on an aluminium foil (1). As a result of the impact, the aluminium foil breaks and the rat’s head makes a rotating movement (2). Immediately upon impact, a spO_2_ sensor is placed on the rat’s hind paw (3). **B)** spO_2_ (mean ± standard error) is reduced at 1 minute post-injury (n=9 rats) compared to isoflurane controls (n=7 rats, p=0.005, Mann-Whitney-test). **C)** NSS scores are significantly decreased at 20 minutes after TBI (n=30 rats, p=0,003, Kruskal-Wallis test), but follow a normal distribution (p=0.29, Anderson-Darling test). At 48h, NSS scores no longer follow a normal distribution (p=0.011, Anderson-Darling test) and two populations can be identified. **D)** Post-mortem structural analysis reveals bruising (arrow) on the skull (bregma indicated with arrowhead), but no cortical structural damage. Hematoma around the brainstem is observed. Scale bar=1 cm. **E)** Total ATPase activity in cortex homogenates is not different two days after TBI (n=4 rats) compared to controls (n=4 rats, p=0.057, Mann-Whitney test), but the contribution of α2/3 isoform ATPase is increased (p=0.029, Mann-Whitney test).

### 3.2 Cortical spreading depolarization causes neurobehavioral decline

Consistent with previous studies (Ritter et al., 2016; Aboghazleh et al., 2021), we recorded CSDs and associated spreading depression of activity in both hemispheres immediately following TBI (Fig. 2A). In a few animals we recorded seizure activity, which occurred either before or after CSD (Fig. 2B). Both CSDs and seizures occurred while spO_2_ was still recovering (Fig. 2A,B). However, CSD was recorded frequently (n=54 out of 106 impacts, 50.9%), while seizures were rare (n=5, 4.7%) (Fig. 2C). To test whether CSD is associated with altered neurobehavioral outcome (Fig. 1E), we measured neurological severity scores (NSS, see Methods) after TBI-induced or electrically triggered CSD. In the setting of TBI, CSD resulted in significantly lower NSS scores compared to animals that did not experience CSD (Fig. 2D). Similarly, in non-impacted control animals, NSS was lower following electrically triggered CSDs compared to sham-operated controls (Fig. 2E), indicating that the occurrence of CSD may underly early post-traumatic neurobehavioral deficits.

**Figure 2.**
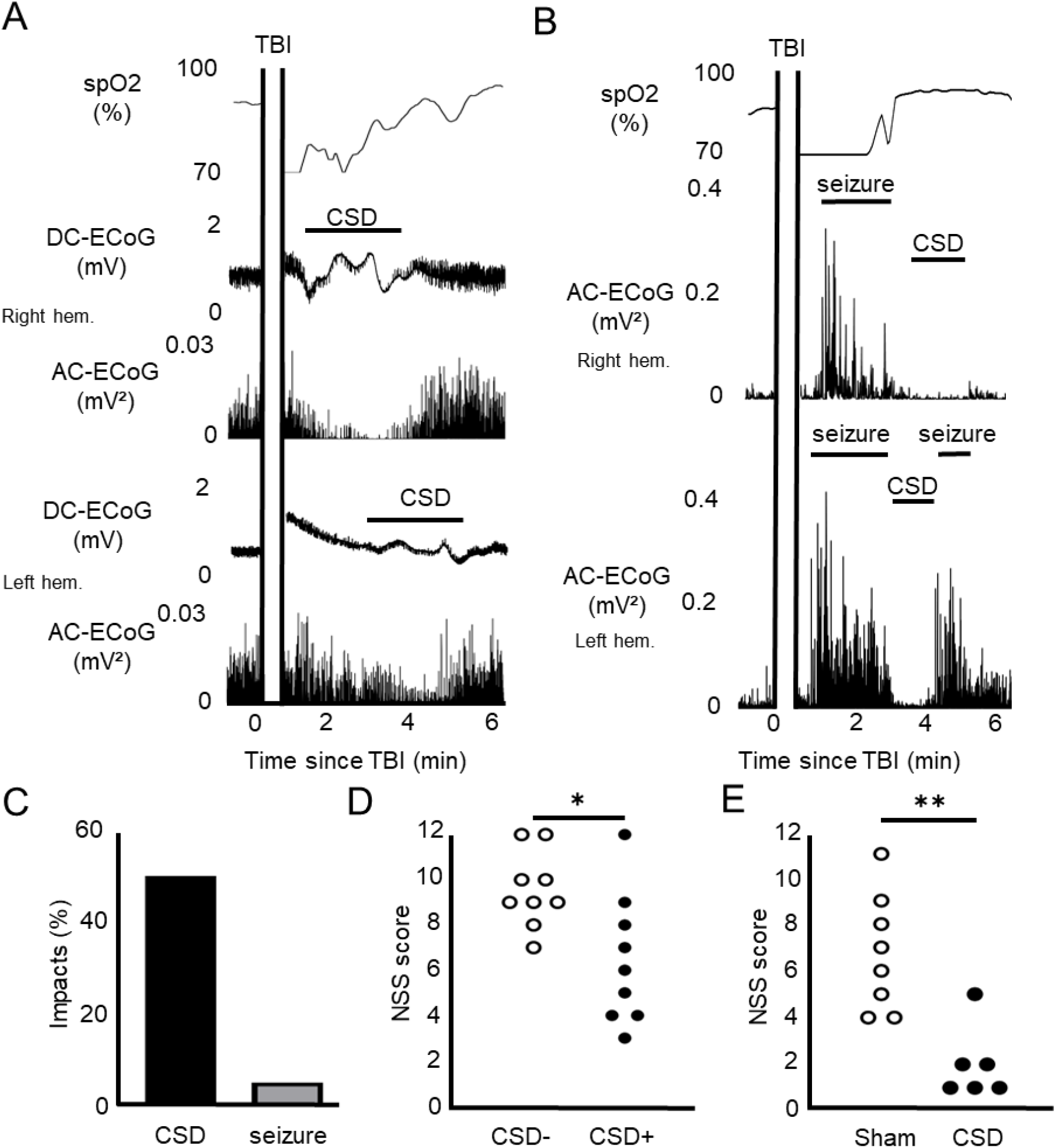
CSD is the most common pathological electrophysiological phenomenon after TBI, linked to behavioral decline. **A)** Representative spO_2_ and ECoG recordings from both hemispheres immediately after TBI with TBI induced CSD. **B)** Representative spO_2_ and ECoG recordings from both hemispheres immediately after TBI induced seizure activity and CSD. **C)** Incidence of CSD after TBI is high (50.9%), while seizures incidence is rare (4.7%). **D)** After head impacts that elicited CSD (n=9), NSS was significantly lower than after head impacts without CSD (n=9, p=0.015, Mann-Whitney test). **E)** Electrically triggered CSD (n=6 rats) did significantly reduce NSS scores at 25 minutes following stimulation (p=0.0016, Mann-Whitney test) compared to sham surgery controls (n=8 rats).

### 3.3 TBI results in vascular dysfunction during and following CSD

Since CSD can be associated with an abnormal vascular response (Dreier et al., 2018; Hinzman et al., 2014), we assessed vascular changes after electrically triggered CSD in sham controls and TBI-injured animals. In TBI rats, suppression of brain activity during CSDs was significantly more pronounced compared to sham controls (Fig. 3A,B). As previously reported, CSDs were associated with a typical vascular response (Ayata and Lauritzen, 2015; Back et al., 1994; Piper et al., 1991) composed of vasodilation with increased local CBF (the hyperemic phase), followed by vasoconstriction associated with decrease in blood flow (an “oligemic phase”). Notably, while the no differences were found in the vascular response to CSD during the hyperemic phase, vasoconstriction was more pronounced in TBI-exposed compared with sham controls (Fig. 3C-F). Despite more pronounced post-CSD oligemia, no differences in brain tissue oxygen partial pressure were measured between TBI injured animals and controls (Fig. 3G,H). Both TBI-injured and sham controls showed an early decrease followed by an increase in available oxygen (Fig. 3G). This observation likely reflects an increase in oxygen consumption and an increase in oxygen delivery through vasodilation (Fig. 3C).

**Figure 3.**
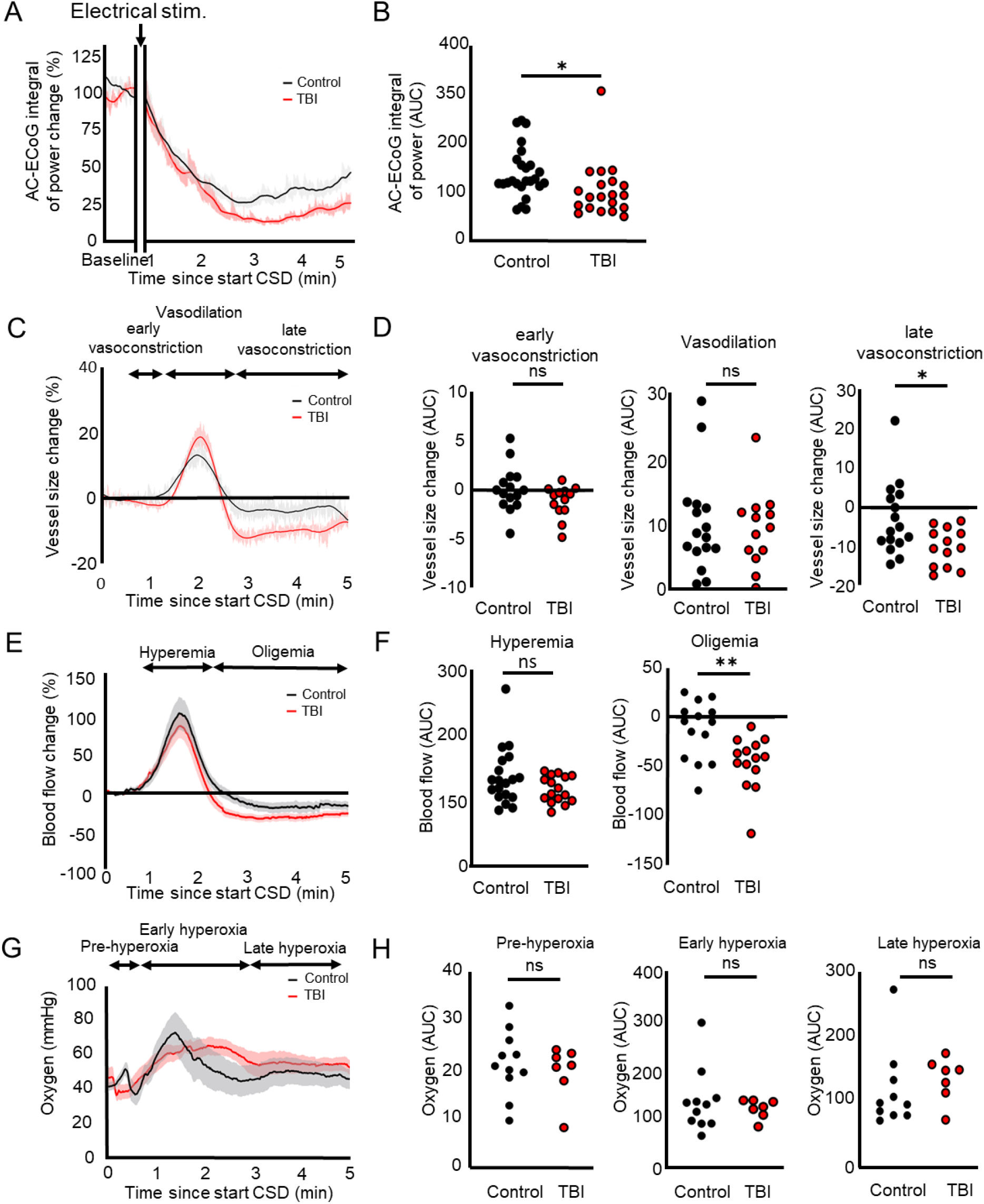
Vascular dysfunction after CSD is worsened in TBI animals. **A)** The alternating current ECoG demonstrates reduction of cortical activity in the minutes following CSD in uninjured (n=25 rats) and TBI (n=20 rats) animals. **B)** Depression of cortical activity is significantly different between uninjured and TBI animals (p=0.032, unpaired t-test). **C)** The vascular diameter change during CSD demonstrates an initial vasoconstriction phase, a vasodilation phase, followed by a second more profound vasoconstriction phase in both uninjured (n=16 rats) and TBI animals (n=13 rats). **D)** No difference in vessel size change is measured between uninjured and TBI animals during the first vasoconstriction phase (p=0.11, t-test), or the vasodilation phase (p=0.76, t-test), but TBI animals show significantly worsened vasoconstriction (p=0.017, t-test). **E)** Blood flow measurements during CSD demonstrate a hyperemic and oligemic phase in both uninjured (n=17 rats) and TBI animals (n=17 rats). **F)** No change in blood flow is measured during the vasodilation phase (p=0.08, t-test), but TBI animals show a worsened oligemic response (p=0.004, t-test). **G)** Changes in cortical oxygen pressure during CSD in untreated (n=11 rats) and TBI (n=9 rats) rats. **H)** No significant difference between TBI and sham control animals was measured in the pre-hyperoxia phase (p=0.79, Mann-Whitney test), the hyperoxia phase (p>0.99, Mann-Whitney test), or the post-hyperoxia phase (p=0.23, Mann-Whitney test).

### 3.4 CSD results in increased mitochondrial ROS

Despite the vasoconstriction and reduced blood flow following CSD, no hypoxia was measured. In fact, in 4 out of 9 TBI animals, oxygen pressure rose after each consecutive CSD (Fig. 4A). Additionally, the maximum oxygen tension during CSD was reached later in TBI animals compared to sham controls (Fig. 4B). Given that oxygen delivery was not impaired in TBI animals (no CBF differences during the hypermic phase), we hypothesized that oxygen consumption by mitochondria is impaired in these TBI animals. We therefore assessed mitochondrial function during CSD using fluorescent dyes. Rhod2-AM was used as an indicator of intramitochondrial calcium (Pozzan and Rudolf, 2009), and MitoSOX as an indicator of mitochondrial ROS production (Mayer et al., 2015; Robinson et al., 2006). Rhod2-AM fluorescence increased during CSD (Fig. 4C). MitoSOX fluorescence similarly increased during CSDs, followed by a decrease in intensity (Fig. 4C). Interestingly, following five consecutive CSDs, a prolonged increase in mitoSOX fluorescent intensity was observed (Fig. 4D) mainly nearby large vessels. Cells within 100 µm from a vessel showed a stronger increase in mitoSOX fluorescence compared to cells located more distally from the vasculature (Fig. 4D,E).

**Figure 4.**
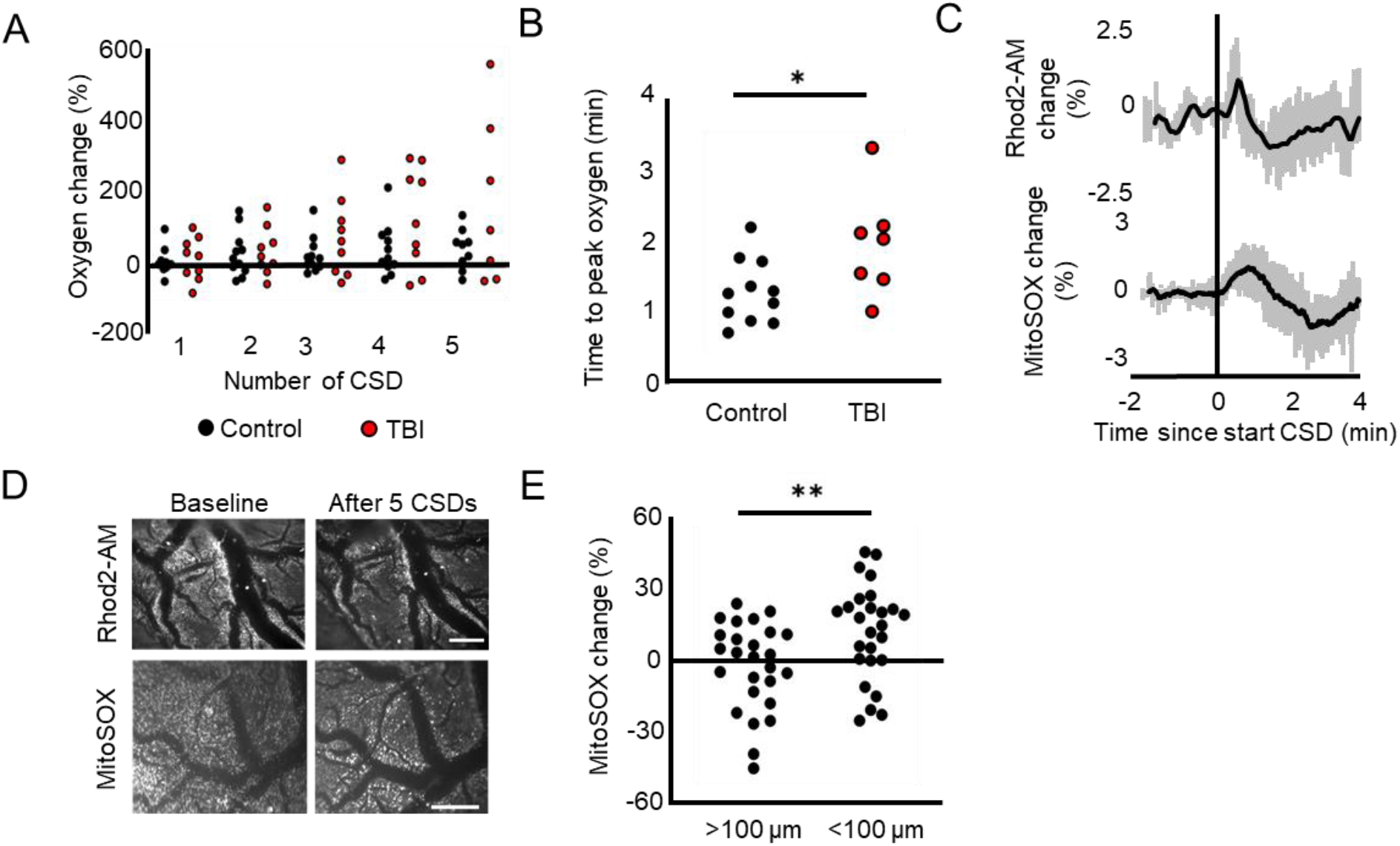
Mitochondrial ROS and calcium during CSD. **A)** The change in oxygen availability after 5 consecutive CSDs is not different between uninjured and TBI animals, although few TBI animals (4 out of 9) show remarkable increase in oxygen pressure. **B)** The time to reach peak oxygen partial pressure during the hyperemic phase of CSD is delayed in TBI animals compared to sham controls (p=0.035, Mann-Whitney test) **C)** During CSD, a spike in Rhod2-AM (n=18 rats) and mitoSOX (n=26 rats) fluorescent intensity is measured (means ± standard error). **D)** A decrease in Rhod2-AM fluorescence, but persistent increase in mitoSOX fluorescence is measured after 5 consecutive CSD, especially around dilating vessels. Scale bar = 1 mm. **E)** The increase in mitoSOX fluorescence is stronger around large vessels compared to cells further away form dilating vessels (n=26 rats, p=0.0016, Paired t-test).

### 3.5 Traumatic brain injury and CSD result in vascular morphological changes

Given the observed increase in oxidative stress in response to CSD, we hypothesised that the vascular dysfunction in TBI animals (i.e. worsened vasoconstriction and hypoperfusion, Fig. 3C-F) is due to mitochondrial dysfunction. Therefore we assessed TBI-induced vascular changes using electron micrographs from cortical sections of rat brains two days after a moderate head impact. Following TBI, cortical vessels appeared contracted with reduced circularity compared to controls (Fig. 5A,B). Similarly, we observed vessels with reduced circularity after triggered CSD in sham controls (Fig. 5C). Since astrocytes and pericytes are two main cell types regulating vascular tone, we also measured size of astrocyte endfeet and pericytes. Astrocyte endfeet (Fig. 5D) did not differ in size in TBI animals compared to controls (Fig. 5E). No differences in astrocyte endfeet size were measured after triggered CSDs compared to sham surgery controls (Fig. 5F). Average pericyte size was also not different in TBI-impacted brains, but was higher in brains in which CSDs were triggered (Fig. 5G-I). Finally, we measured tight junction diameter (Fig. 5J). No differences in tight junction diameter were found after TBI (Fig. 5K), nor after CSDs (Fig. 5L).

**Figure 5.**
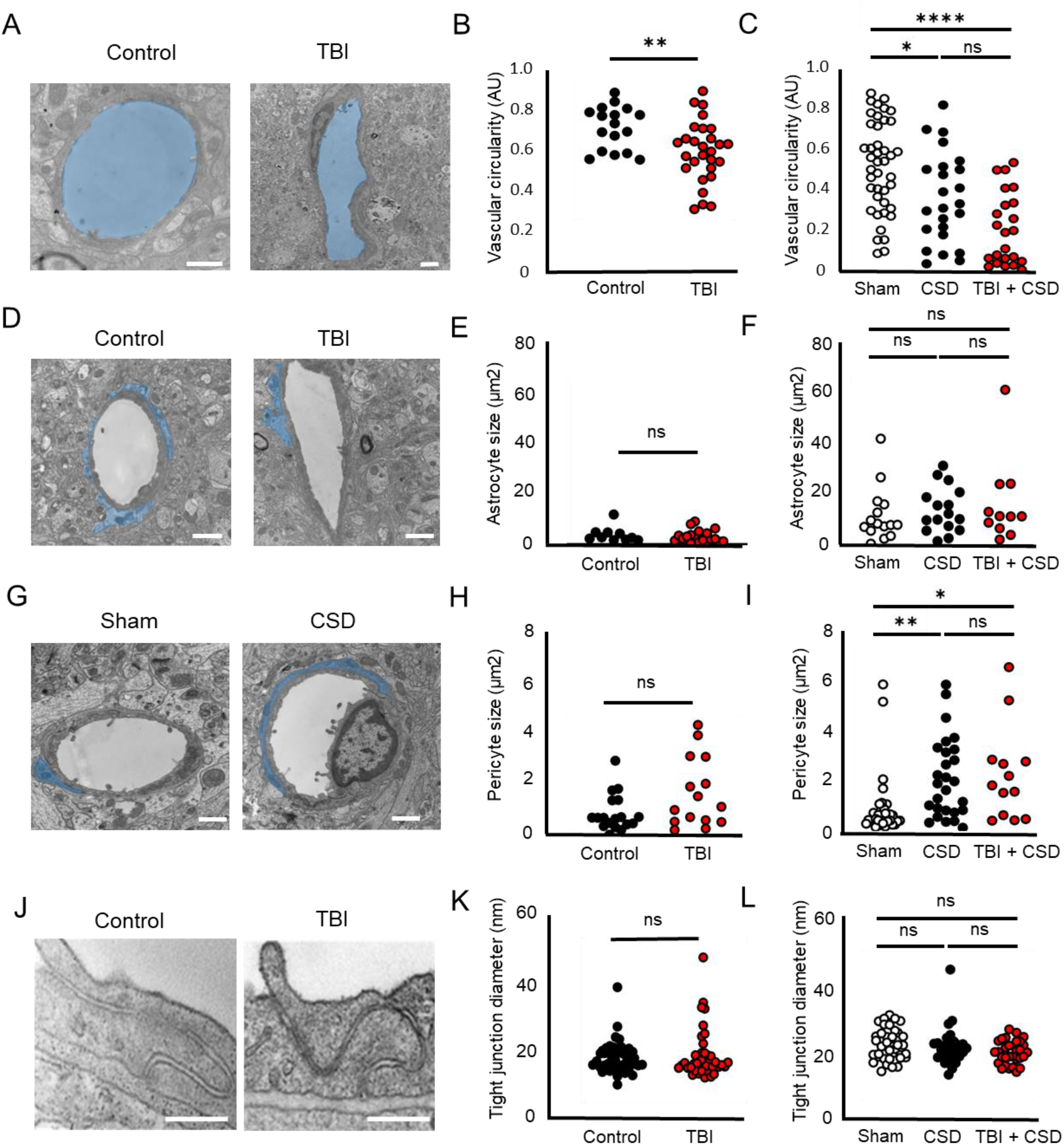
Vascular morphological changes following TBI and CSD. **A)** Representative electron micrographs of a cortical vessel in healthy rat brain (n=4 rats) and after TBI (n=4 rats) with lumen indicated in blue. Scale bar=1 µm. **B)** The vessel circularity is significantly reduced in TBI impacted animals (n=28 vessels, p=0.007, unpaired t-test) compared to healthy isoflurane treated control animals (n=17 vessels). **C)** Vessel circularity is significantly reduced after CSD (n=24 vessels from 5 rats, p=0.045, Kruskal-Wallis test) and after TBI plus triggered CSD (n=23 vessels from 4 rats, p<0.0001, Kruskal-Wallis test) compared to sham surgery controls (n=42 vessels from 4 rats). **D)** Representative electron micrographs of healthy rat brain (left) and after TBI + CSD (right, n=4 rats) with astrocyte endfeet indicated in blue. Scale bar=1 µm. **E)** Astrocyte endfeet size is the same between healthy (n=10 cells) and TBI animals (n=19 cells, p=0.38, Mann-Whitney test). **F)** Astrocyte endfoot size is not different between sham controls (n=15 cells) and after CSD (n=16 cells, p=0.96, Kruskal-Wallis test) or after TBI plus triggered CSD (n=11 cells, p>0.99, Kruskal-Wallis test). **G)** Representative electron micrographs of healthy rat brain (left) and after CSD (right) with pericytes indicated in blue. Scale bar=1 µm. **H)** Pericyte size is the same between healthy (n=18) and TBI animals (n=15 cells, p=0.08, Mann-Whitney test). **I)** Pericyte size is significantly larger after CSD (n=27 cells, p=0.0011, Kruskal-Wallis test) and after TBI plus triggered CSD (n=13 cells, p=0.01, Kruskal-Wallis test) compared to sham surgery controls (n=31 cells). **J)** Representative electron micrographs of tight junctions in healthy rat brain (left) and after TBI (right). Scale bar=200 nm. **K)** Tight junction diameter is the same between healthy (n=47 tight junctions) and TBI animals (n=41 tight junctions, p=0.58, unpaired t-test). **L)** Tight junction diameter is the same between sham controls (n=38 tight junctions) and after CSD (n=30 tight junctions, p>0.99, Kruskal-Wallis test) or after TBI plus triggered CSD (n=30 tight junctions, p=0.19, Kruskal-Wallis test).

### 3.6 Traumatic brain injury and CSD result in mitochondrial swelling and cristae damage in astrocytes, pericytes and endothelial cells

Since we measured mitochondrial ROS increases around large vessels upon triggered CSD, mitochondrial morphology was examined (Fig. 6A). The size of mitochondria was greater in pericytes and neurons from TBI-exposed brains compared with sham controls (Fig. 6 A,B), suggesting swelling. No change in mitochondrial size was measured in astrocytes, endothelial cells or myelinated axons (Fig. 6B). We also noted evidence for mitochondrial cristae damage (Kirov et al., 2020) in pericytes, astrocytes and neurons (except in myelinated parts of the axon) from TBI-brains but not in endothelial cells (Fig. 6A-C).

Triggering CSD in healthy control rats resulted in morphologically similar astrocytic mitochondrial changes (i.e. swelling and cristae damage) compared to changes occurring following TBI, supporting the notion that post-traumatic CSDs underlie the observed differences (Fig. 6D,E). In contrast, mitochondria morphology in pericytes and endothelial cells were not affected by electrically triggered CSDs (Fig. 6D,E). Triggering CSDs in TBI brains resulted in mitochondrial cristae damage in endothelial cells as well (Fig. 6E), suggesting that TBI may cause endothelial cells to become vulnerable to additional metabolic challenges, such as CSD. Triggering CSD in TBI brains resulted in mitochondrial cristae damage in endothelial cells as well (Fig. 6E), suggesting that TBI caused endothelial cells to become vulnerable for additional metabolic challenges, such as CSD. Injury to mitochondria in neuronal somata (but not in myelinated portions of the axon) was observed after CSD in both controls and TBI-exposed brains (Fig. 6E).

**Figure 6.**
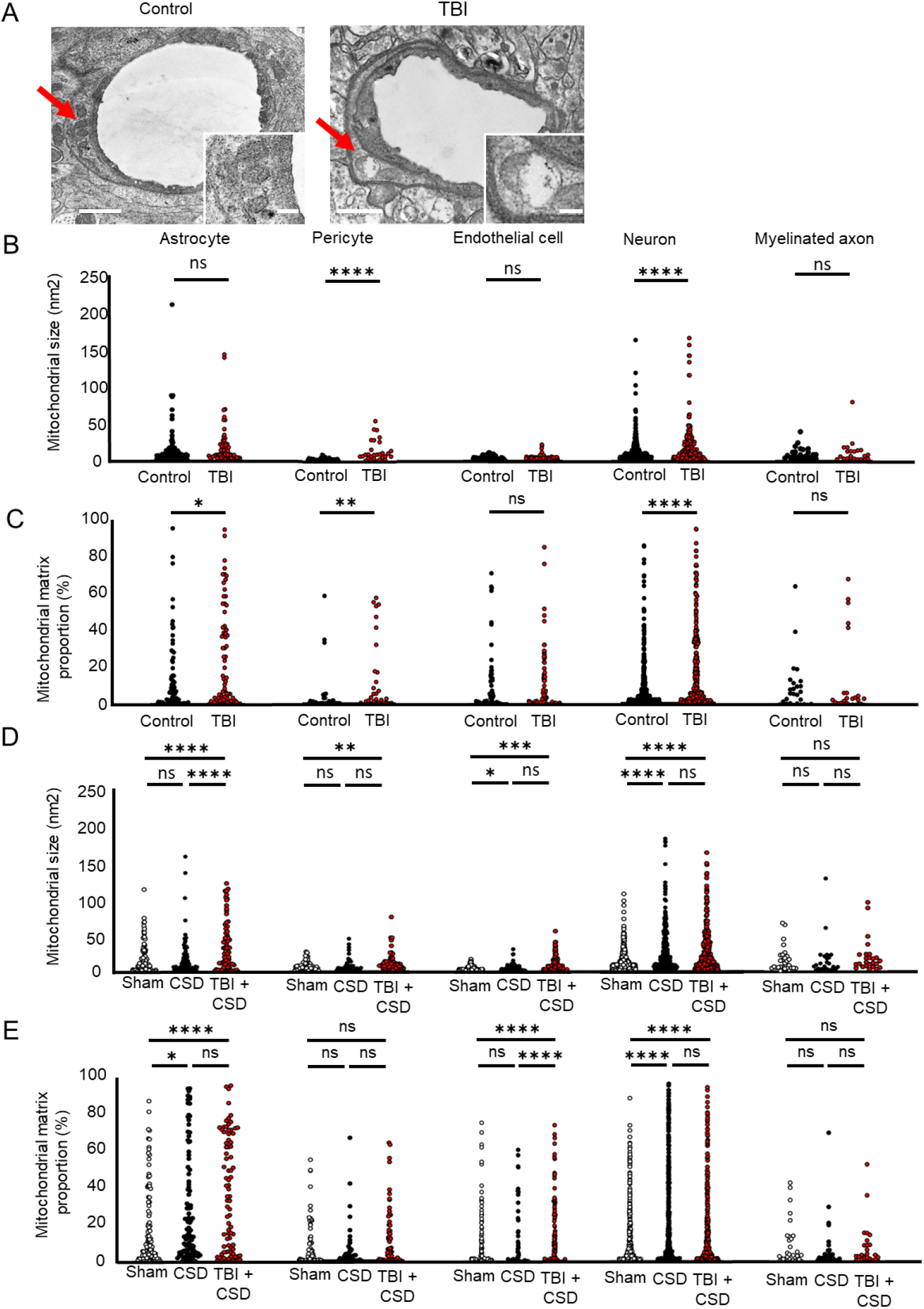
Mitochondrial morphological changes in vascular cells following TBI and CSD. **A)** Representative electron micrographs of two vessels in healthy rat brain (left) and after TBI (right). Pericytes indicated by arrows. Scale bar=1 µm. Representative pericyte mitochondria are shown in the inserts on the bottom right. Scale bar of insert=200 nm. **B)** Mitochondria in pericytes (n=62 mitochondria, p<0.001) and neurons (n=785 mitochondria, p=<0.001) are significantly larger in size in TBI brains than in healthy brains, but not in astrocyte (n=163 mitochondria, p=0.23) and endothelial cell mitochondria (n=214 mitochondria, p=0.59), and mitochondria in myelinated axons (n=54 mitochondria, p=0.75). Significance tested via Mann-Whitney tests. **C)** Mitochondrial show cristae damage after TBI in astrocytes (p=0.014), in pericytes (p=0.002), and in neurons (p<0.0001), but not in endothelial cells (p=0.65) or in myelinated axons (p=0.28). Significance tested via Mann-Whitney tests. **D)** Mitochondria in endothelial cells (n=452 mitochondria, p=0.02) and neurons (n=1454 mitochondria, p<0.001) are significantly larger in size after CSD compared to sham surgery controls, whereas mitochondria in astrocytes (n=296 mitochondria, p>0.99), pericytes (n=171 mitochondria, p=0.67), and myelinated axons (n=85 mitochondria, p>0.99) are not. After triggered CSD in TBI animals, astrocyte (p=0.001), pericyte (p=0.003), endothelial cell (p=0.0003), and neuronal (p<0.0001) mitochondria are larger than sham controls, but not myelinated axons (p=0.93). Significance tested via Kruskal-Wallis tests. **E)** Mitochondria show cristae damage after CSD in astrocytes (p=0.01) and in neurons (p<0.0001), but not in pericytes (p>0.99), endothelial cells (p>0.99) or in myelinated axons (p>0.99). After TBI plus triggered CSD, mitochondria show cristae damage in astrocytes (p<0.0001), endothelial cells (p=0.0002), and neurons (p<0.0001), but not in pericytes (p=0.25) or myelinated axons (p>0.99). Significance tested via Kruskal-Wallis tests.

### 3.7 TBI results in vascular dysfunction following seizures

In addition to CSD, seizures were also reported as an acute brain response to injury (Fig. 2C, Ritter et al., 2016). We therefore measured vascular and mitochondrial functional changes in response to seizures induced by direct cortical application of 4-AP in TBI- and sham-control animals. Recurrent seizures were recorded using silver wire electrode ECoG (Fig. 5Aa). Similar to CSD, during seizures, increased CBF (Fig. 7Ab), brain oxygen partial pressure (Fig. 7Ac) and mitochondrial ROS production (Fig. 7Ad) were observed. A transient drop in oxygen pressure at the beginning of seizures, was often observed (27 out of 31 seizures), both in TBI-exposed animals and controls (Fig. 5Ac). In TBI-exposed animals, 4-AP application resulted in a higher frequency of recurrent seizures (Fig. 5B), with a less pronounced increase in local CBF compared with controls (Fig. 5C-D).

**Figure 7.**
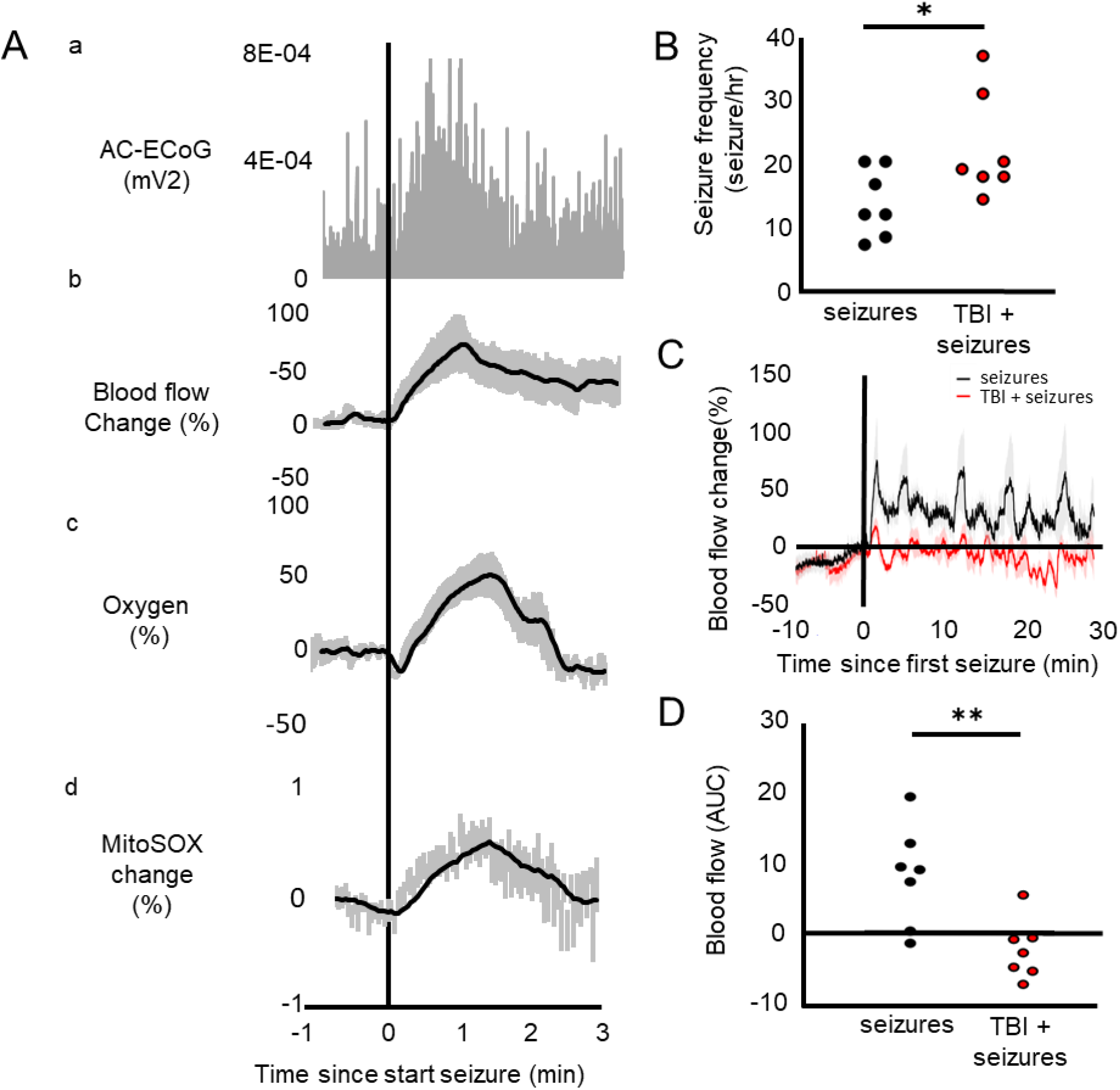
The vascular and mitochondrial response to seizures in TBI animals. **A)** Seizure activity is identified by neuronal hyperactivity in the AC-ECoG recording (a). During seizures, blood flow increases (b, n=7 rats), as well as brain oxygen partial pressure (c, n=34 seizures in 2 rats). MitoSOX fluorescence also increases during seizure activity (d, n=31 seizures in 10 rats). **B)** In TBI animals (n=7 rats), 4-AP induced seizure frequency is significantly higher than in uninjured animals (n=7 rats, p=0.04, t-test). **C)** Changes in blood flow during seizure activity in uninjured (n=7 rats) and TBI (n=7 rats) animals. **D)** Cortical blood flow during seizure activity is significantly lower in TBI compared to uninjured animals (p=0.006, t-test).

## 4. Discussion

Secondary brain damage occurring in the hours to days after head trauma is a promising target for early post-traumatic intervention. However, the mechanisms underlying secondary brain damage are poorly understood. Here, we demonstrate mitochondrial damage after head impact in NVU cells. We also show that electrically triggered CSD cause similar mitochondrial damage in the NVU and increased mitochondrial ROS nearby large vessels. Together, this suggests that mitochondrial dysfunction underlies abnormal vascular response to increased metabolic demand, such as that during CSD and seizures. Mitochondrial and vascular dysfunction during CSD could therefore underly poor TBI outcome.

We demonstrate physiological, structural, and neurobehavioral changes in our moderate TBI model. These changes include immediate low spO_2_, epidural hematoma, and increased α2/3 ATPase activity. NSS scores were significantly reduced at 20 minutes after head impact. At 48 hours after impact, we observed a bimodal NSS distribution, suggesting a pathological event that occurs in some animals, but not others, and which affects behavioral outcome. We hypothesized that pathological electrophysiological events after TBI could explain these behavioral outcomes and found that CSD was occurring frequently (∼50% of impacts), while seizures were rare. Electrically elicited CSD could also significantly reduce NSS. This finding is in accordance with previous results from our lab showing that TBI induced CSD in mild TBI is associated with a longer time to regain locomotion (Parker et al., 2022).

To understand the mechanism through which CSD can cause behavioral decline, we electrically triggered CSD and assessed the functional changes in the brain *in vivo.* We first studied the effect of CSD on the uninjured brain to assess its contribution on the acute, primary brain damage. As previously described (Ayata and Lauritzen, 2015), triggered CSD was associated with initial hypoperfusion (phase I), followed by peak hyperemia (phase II), late hyperemia (phase III) and post-CSD oligemia (phase IV). While hyperemia lasts only a couple of minutes, the oligemic phase may last for hours (Ayata and Lauritzen, 2015). Our results show a drop in oxygen pressure either immediately before or early during vasodilation, which can be the result of early hypoperfusion (phase I), but could also be due to increased consumption of oxygen during oxidative phosphorylation by mitochondria. During TBI induced CSD, when blood oxygenation levels are low (Fig. 1C), mitochondria’s oxygen consumption need might not be met by the vasculature’s supply, which could lead to inverse neurovascular coupling (Prager et al., 2019) and mitochondrial dysfunction (Almeida et al., 2002; Varela et al., 2016). We then measured a rise in oxygen during CSD, which is in contrast with previous research that report severe hypoxia during spreading depolarization (Dreier, 2011; Takano et al., 2007). The difference could be due to the constant supply of oxygen during isoflurane anaesthesia in our experimental setup.

We further report a change in mitochondrial function in response to triggered CSD. Production of ROS increased during CSD, pointing to increase in oxidative phosphorylation. The increase in mitochondrial ROS was followed by a decrease, which could reflect cytotoxic edema, characteristic to CSDs (Dreier et al., 2018; Kirov et al., 2020). The excess of ion and water influx will also affect mitochondria and result in swelling of the mitochondria (Zhou et al., 2010) which may explain a relative decrease (dilutional effect) in mitoSOX and Rhod2-AM fluorescence intensities. Following repetitive CSD, we found that mitochondrial ROS production remained high, most prominent in the vicinity of dilating vessels, consistent with the role of mitochondrial ROS in the regulation of vascular tone (Totzeck et al., 2014). This finding is in accordance with a previous study reporting increased NADH fluorescence, a redox state indicator, in the proximity of vasculature following CSDs (Takano et al., 2007).

CSD, oligemia, and mitochondrial ROS are thought to contribute to secondary brain damage following injury (Dreier et al., 2006; Hartings et al., 2020; Hiebert et al., 2015; Toth et al., 2016; Verweij et al., 2007). Therefore, to test whether the injured brain respond differently to metabolic challenges, we electrically triggered CSD, or pharmacologically induced seizures two days following TBI. This way, we assessed the effect of CSD in delayed, secondary brain damage. We report vascular dysfunction after CSD or seizures in injured animals, with stronger vasoconstriction and worsened hypoperfusion. Changes in brain oxygen levels during CSD were not significantly different between TBI and healthy animals, but 4 out of 9 TBI animals showed a remarkable increase in brain oxygen. Such increase in oxygen has been previously attributed to low neuronal metabolism during burst suppression (Berndt et al., 2021). Alternatively, impaired mitochondrial oxygen consumption in these injured brains, without affected oxygen delivery, may result in abnormally high oxygen partial pressure (Verweij et al., 2000). Similarly, regional CBF elevation in response to seizures was lacking in TBI animals, demonstrating that injured brain damage results in impaired regulation of regional CBF during metabolic challenges (Balança et al., 2017). Additionally, seizure frequency was higher in TBI animals. TBI induced mitochondrial damage in neurons could underly the lowered seizure threshold, since Styr et al. demonstrated mitochondria’s role in regulating excitability thresholds through calcium buffering (Styr et al., 2019).

Electron microscopy images further supported vascular and mitochondrial damage after TBI and triggered CSDs. Vascular injury manifested as reduced vessel circularity or enlarged pericytes, which is in accordance with previous studies (Ichkova et al., 2020; Lin et al., 2022; Parker et al., 2022; Prager et al., 2019; Rafols et al., 2007) and may underlie TBI-induced dysfunction of the blood-brain barrier (BBB). Mitochondrial damage was observed after TBI and triggered CSDs, which has also been demonstrated in previous studies (Verweij et al., 2000; Vink et al., 1990). However, to our knowledge, we demonstrate for the first time that neurons, astrocytes, and pericytes are more prone to mitochondrial damage than endothelial cells. A potential reason is that the mitochondrial content is relatively high in endothelial cells (Oldendorf et al., 1977) and therefore their capacity to mitigate pathological molecular mechanisms (i.g. calcium buffering) after head impacts could be better.

We observed mitochondrial damage in both pericytes and astrocytes, two cell types critically involved in regulating local CBF. Neurovascular uncoupling and reduced oxygen have been previously reported in pericyte deficient mice (Kisler et al., 2017), further supporting the notion that mitochondrial damage in these cells may underlie impaired vascular response. Indeed, mitochondrial dysfunction in pericytes reduces contractile capability in diabetic rats (Liu et al., 2021). Since pericytes might not play a role in vasodilation (Fernández-Klett et al., 2010), this could explain why no changes in vasodilation and hyperemia were detected in TBI animals, but vasoconstriction and oligemia was worsened. Astrocytes regulate local CBF via their endfeet and cytosolic calcium concentrations in astrocyte endfeet have been shown to regulate vascular shape (Gordon et al., 2008; Haidey and Gordon, 2021). Although we did not observe astrocytic swelling, which was previously shown to alter vascular tone (Bullock et al., 1991), functional astrocytic changes might underly vascular dysfunction. We found mitochondrial damage in astrocytic endfeet, which may impair calcium buffering, resulting in abnormal calcium signalling, which has been demonstrated to affect neuronal function (Shigetomi et al., 2019) and CBF (Filosa et al., 2004; Iadecola and Nedergaard, 2007; Zonta et al., 2003). Following TBI and triggered CSD, endothelial cell mitochondria were also damaged. Endothelial mitochondrial damage potentially underlies BBB leakage after CSD in animals pre-exposed to TBI (Parker et al., 2022). Parker et al. found no BBB leakage after CSD in healthy animals, which is in accordance with our observation that endothelial cell mitochondria lack cristae damage. Impaired integrity of mitochondrial membranes in juxtavascular cells was also previously shown to underly BBB opening during seizures (Prager et al., 2019).

In our study, we focussed on the upper cortical regions to match the *in vivo* observations via the cranial window with the *postmortem* analysis. In upper cortical regions, myelination is relatively sparce (van Tilborg et al., 2017) thus only few mitochondria in myelinated axons were observed. Nonetheless, these mitochondria seemed to be protected from CSD-induced swelling and cristae damage. This observation is consistent with the notion that CSD does not propagate into the myelinated axons (Dreier et al., 2018; Merkler et al., 2009)

In conclusion, our study demonstrates that pathological electrophysiological events acutely after head impact can aggravate secondary brain damage. CSD is linked to increased mitochondrial ROS and mitochondrial damage in astrocytes and pericytes. This may cause secondary brain damage, which could manifest as abnormal neurovascular coupling during metabolic challenges in the days after the initial injury. This CSD-induced dysfunction of the vasculature may also explain poor outcome in some TBI patients, while others recover fully. Therefore, CSD and CSD triggered secondary brain damage are promising intervention targets for future studies.

## Abbreviations

4-AP: 4-aminopyridine
AC: alternative current
aCSF: artificial cerebrospinal fluid
ATP: adenosine triphosphate
CBF: cerebral blood flow
CSD: cortical spreading depolarization
DC: direct current
ECoG: electrocorticography
LDF: laser doppler flowmetry
NSS: neurological severity score
ROS: reactive oxygen species
SpO_2_: peripheral oxygen saturation
TBI: traumatic brain injury

## CRediT authorship contribution statement

**Gerben van Hameren:** Conceptualization, Methodology, Formal analysis, Investigation, Writing-Original draft, Writing-Review & Editing, Visualization, Funding Acquisition **Jamil Muradov:** Methodology, Formal analysis, Investigation, Writing-Review & Editing **Anna Minarik** Formal analysis, Investigation, Writing–Review & Editing **Refat Aboghazleh** Methodology, Invstigation **Sophie Orr** Formal analysis, Investigation **Shayna Cort** Methodology, Formal analysis, Investigation **Keiran Andrews** Formal analysis, Investigation **Caitlin McKenna** Investigation **Nga Thy Pham** Investigation **Mark Mclean** Investigation, Writing–Review & Editing **Alon Friedman** Conceptualization, Writing–Review & Editing, Supervision, Funding Acquisition

## Funding sources

This work was supported by the Canadian Institute for Health Research PJT 148896 and by the U.S. Army Medical Research Acquisition Activity Office, 820 Chandler Street, Fort Detrick MD 21702-5014, as the awarding and administering acquisition office of the Assistant Secretary of Defense for Health Affairs, through the Epilepsy Research Program, under Award No. W81XWH-17-1-0684. Opinions, interpretations, conclusions, and recommendations are those of the author, and are not necessarily endorsed by the Department of Defense. In addition, the study was supported by Mitacs Accelerate (IT13603) and by Citizens United for Research in Epilepsy, doing business as CURE Epilepsy (CURE Epilepsy).

## Declaration of competing interests

The authors declare no conflict of interest.

## Acknowledgement

The authors thank Kathleen Murphy, Mary Ann Trevors and Jim Kukurin for technical assistance.

## Notes

### Competing Interest Statement

The authors have declared no competing interest.

